# Non-invasive Imaging Techniques for Monitoring Cellular Response to Treatment in Stable Vitiligo

**DOI:** 10.1101/2023.08.15.553419

**Authors:** Jessica Shiu, Griffin Lentsch, Christopher M. Polleys, Pezhman Mobasher, Marissa Ericson, Irene Georgakoudi, Anand K Ganesan, Mihaela Balu

## Abstract

Punch grafting procedures, where small pieces of normal skin are transplanted into stable vitiligo patches, results in repigmentation in only half of patients treated, yet the factors that determine whether a patient responds to treatment or not are still unknown. Reflectance confocal microscopy (RCM) is adept at visualizing melanocyte migration and epidermal changes over large areas while multiphoton microscopy (MPM) can capture metabolic changes in keratinocytes. With the overall goal of identifying optical biomarkers for early treatment response, we followed 12 vitiligo lesions undergoing punch grafting. Dendritic melanocytes adjacent to the graft site were observed before clinical evidence of repigmentation in treatment responsive patients but not in treatment non-responsive patients, suggesting that the early visualization of melanocytes is indicative of a therapeutic response. Keratinocyte metabolic changes in vitiligo skin adjacent to the graft site also correlated with treatment response, indicating that a keratinocyte microenvironment that more closely resembles normal skin is more hospitable for migrating melanocytes. Taken together, these studies suggest that successful melanocyte transplantation requires both the introduction of new melanocytes and modulation of the local tissue microenvironment.

## Introduction

Vitiligo, an autoimmune skin disease affecting 0.5% to 2% of the global population, is characterized by the progressive destruction of melanocytes by autoreactive CD8^+^ T cells [1-11]. This chronic condition leads to disfiguring white patches, causing depression, anxiety and social isolation among patients [11-15]. The gold standard treatment for vitiligo is the combination of narrow band UVB (nbUVB) light therapy and topical immune-suppressing treatments [16-20]. Recent advancements, such as the addition of topical Jak inhibitors (JAKi) to nbUVB, have improved responses, particularly in facial areas, although patients still experience partially repigmented skin [21, 22]. Surgical interventions such as punch grafting and non-cultured epidermal suspensions (NCES) are used to repopulate depigmented tissue with melanocytes in stable vitiligo patches but, only about 45-47% of patients achieve satisfactory results with >90% repigmentation respectively [23]. Treatment response varies by patient age, vitiligo site and vitiligo subtype, with segmental vitiligo having the most successful response. Pigment spread is usually evident around 1 month but there is no clear way to identify patients that would respond to therapy [24]. Grafting methods differ in their requirement for destruction of the entire epidermis at the graft recipient site. It is currently unclear how much local factors at the recipient site influence response to grafting therapies.

Reflectance confocal microscopy (RCM) and multiphoton microscopy (MPM) are noninvasive imaging techniques [25-27] that enable cellular-level monitoring of treatment response in skin without the need for labeling [28-30]. RCM has been used to assess vitiligo stability in patients [31] and to characterize vitiligo before and after treatment [32]. MPM offers the additional advantage of molecular contrast, allowing for the detection of cellular metabolic changes through the two-photon excited fluorescence (TPEF) signal emitted from reduced nicotinamide adenine dinucleotide (NADH) [33, 34]. In a recent study, our group employed *in vivo* MPM imaging and single cell RNA-sequencing analysis to identify a metabolically altered population of keratinocytes in vitiligo skin that plays a significant role in controlling vitiligo persistence [35]. Although MPM provides the advantage of label-free molecular contrast for metabolic imaging, the current commercial clinical device based on this technology, has a limitation regarding the imaging area, which is reduced to sub-millimeter scale. To overcome this limitation, MPM imaging can be combined with RCM clinical imaging, which allows for the examination of larger skin areas in the multi-millimeter range [27].

In this study, we employ both RCM and MPM technologies to monitor the treatment response to punch grafting in vitilgo patients. Using RCM, we can visualize whether early detection of melanocyte migration correlates with treatment response. MPM allows the examination of metabolic changes in vitiligo skin locally to determine whether repigmentation occurs more often in skin that has a metabolic profile more akin to normal skin. Here we combine two imaging technologies to examine how melanocyte migration and the local tissue microenvirnoment influence grafting response. Understanding these factors will assist with designing better grafting therapies with improved response rates and developing more quantitative methods to measure melanocyte repopulation after therapy.

## Results

We followed 12 vitiligo lesions in 11 patients to evaluate the effectiveness of treatment by utilizing *in vivo* MPM and RCM imaging across 5 visits (Table 1, Figure 1A). Among these lesions, 5 showed repigmentation, based on visual clinical assessment. The treatment involved punch grafting and nbUVB treatment as described in the *Methods* section. However, not all patients could attend all 5 visits for both imaging modalities. The objective of this study was to assess the ability of MPM and RCM to capture cellular morphological changes associated with keratinocyte pigmentation and melanocyte migration resulting from treatment. Furthermore, we sought to identify optical biomarkers related to melanocyte migration and metabolic changes in keratinocytes that could facilitate early detection of treatment response.

**Table 1.** Patient Characteristics.

| Lesion ID | Age | Sex | Area imaged | Disease Status | Notes |
| --- | --- | --- | --- | --- | --- |
| **1** | 61 | M | Arm | Acrofacial | Clinical Responder |
| **2** | 34 | M | Wrist | Generalized | Missed multiple RCM imaging visits, excluded from RCM analysis |
| **3** | 73 | F | Back | Generalized |  |
| **4** | 45 | F | Hand | Focal | Clinical Responder |
| **5** | 74 | M | Wrist | Acrofacial |  |
| **6** | 58 | M | Leg | Generalized | Clinical Responder |
| **7** | 36 | M | Leg | Generalized |  |
| **8** | 50 | F | Face | Acrofacial | Clinical Responder |
| **9** | 72 | F | Hand | Acrofacial |  |
| **10** | 39 | F | Leg | Generalized | Clinical Responder |
| **11** | 20 | F | Ankle | Generalized |  |
| **12** | 36 | M | Leg | Generalized | Same patient as 7, different lesion |

**Figure 1.**
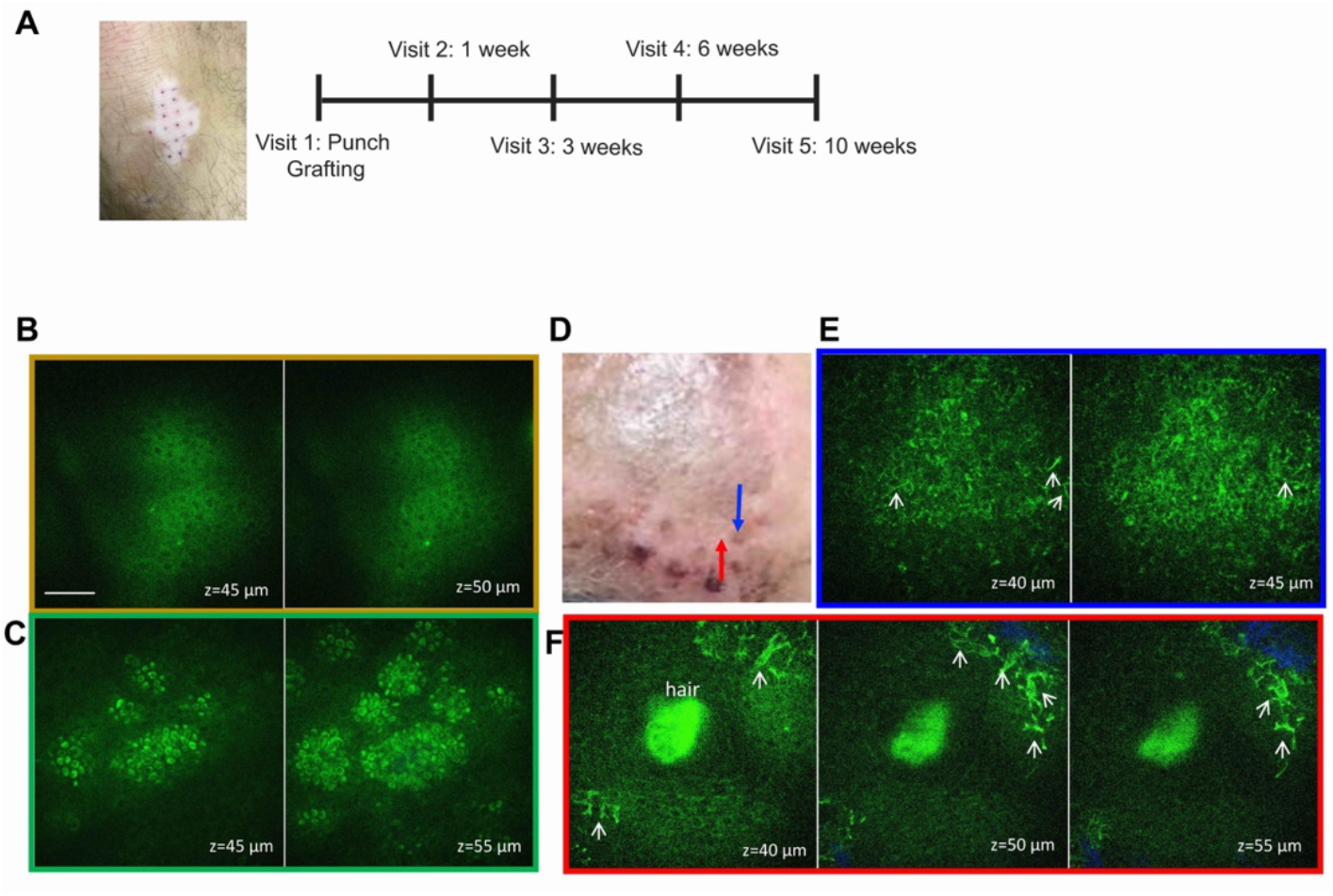
Study design and MPM images of treated vitiligo. (A) Schematic of the study design. Representative en-face MPM images of vitiligo lesional acceptor (B) and nonlesional donor (C) skin acquired *ex vivo* immediately before grafting; (D) clinical image post punch grafting procedure showing the imaged graft (blue arrow) and adjacent imaged location (red arrow). (E) Representative en-face MPM images acquired *in vivo* from the graft indicated in (D) showing evidence of activated melanocytes (arrows). (F) Representative en-face *in vivo* MPM images of the adjacent location to the graft showing migrating melanocytes in the proximity of a hair follicle. Color outline of en-face images in E and F correspond to location indicated by arrows in D.

### MPM captures morphological changes associated with keratinocyte pigmentation and melanocyte migration after punch grafting treatment in vitiligo patients

Figure 1 B-F presents representative clinical and MPM images acquired *ex vivo* and *in vivo* from both the recipient and donor sites in one of the patients pre- and post-treatment. Prior to treatment, MPM imaging revealed the absence of melanocytes and melanin in the recipient site skin (Figure 1B) and the presence of melanin in keratinocytes in the donor site skin (Figure 1C). During the treatment period, the same graft site and an adjacent area were imaged *in vivo* by MPM (Figure 1D). Notably, imaging of the graft site 1 week after procedure revealed the presence of pigmentation (Figure 1E), while large dendritic melanocytes were visualized in areas surrounding the graft site (Figure 1F). Although these images highlight MPM’s capability to visualize the repigmentation process in vitiligo, the limited field of view of the MPM device employed in this study restricted its ability to monitor and quantify the migration of melanocytes throughout the treatment duration.

### RCM can identify optical biomarkers associated with melanocyte migration to facilitate early detection of treatment response in vitiligo patients

The clinical RCM device utilized in this study provided images covering a multi-millimeter range area, allowing us to image the same graft at each visit. This facilitated the accurate assessment of the melanocyte migration following the treatment of vitiligo lesions. We conducted a quantitative analysis based on the RCM images obtained from the same graft and its surrounding area throughout the treatment period for each patient. The optical biomarkers used to quantify melanocyte migration as a response to treatment were the density of melanocytes (number of melanocytes per unit area) and the distance of melanocytes from the graft.

Activated melanocytes featuring bipolar and/or stellate dendrites [36, 37] were observed in all patients showing clinical repigmentation, as early as 3 weeks in two patients and 6 weeks in most patients. Generally, the presence of activated melanocytes preceded clinical evidence of repigmentation. Due to participant dropout for some of the imaging visits, the analysis had to be adjusted accordingly (see methods and Table 1). The number of melanocytes appeared to increase with time. Conversely, no activated melanocytes were detected in vitiligo patients who did not respond to punch grafting treatment (Figure 2A), indicating an association between the presence of activated melanocytes and treatment response.

**Figure 2.**
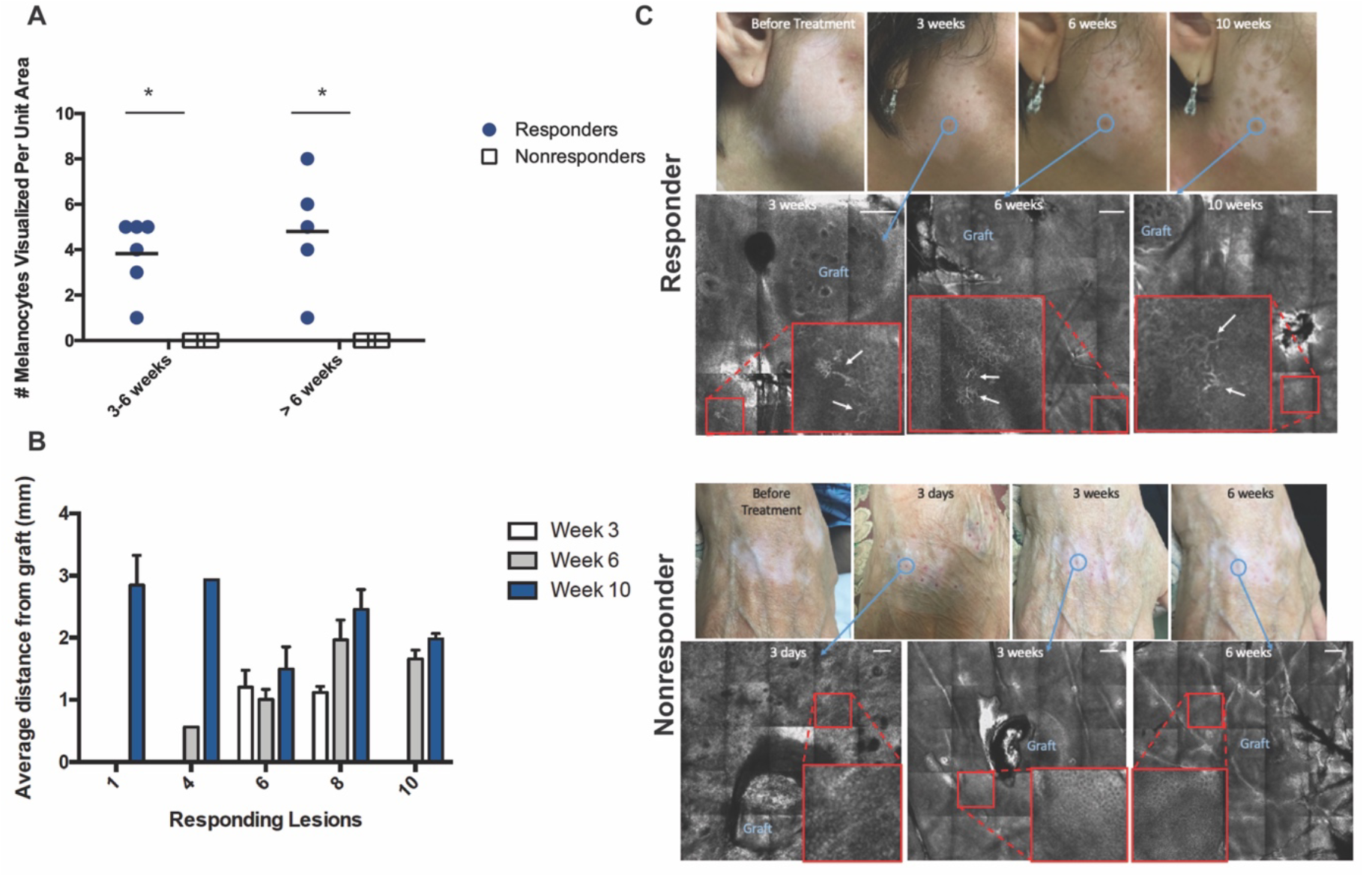
RCM images of skin after punch graft treatment and melanocyte quantification. **(A)** Number of activated melanocytes visualized per unit area in treatment responders and nonresponders at different visits. Each data point represents number of activated melanocytes visualized in a lesion per unit area per visit. * = t-test p-value < 0.05 **(B)** Average distance per melanocyte from punch graft measured in mm in treatment responders. **(C)** Representative clinical and RCM images acquired from the treated vitiligo skin of treatment responders (top) and nonresponders. Blue circle depicts clinical area that was imaged over time and the corresponding RCM image. Enlarged image of area of interest is shown in large red squares with white arrow indicating activated melanocytes. Scale bars are 200 μm.

Analysis of the RCM images obtained at different time points also revealed that in the majority of treatment-responsive patients, the distance of melanocytes from the graft increased over time (Figure 2B), suggesting melanocyte migration. Importantly, no patients in whom melanocytes were imaged showed subsequently disappearance of melanocytes at a later time point.

### MPM can provide optical biomarkers associated with metabolic changes in keratinocytes to facilitate early detection of treatment response in vitiligo

The MPM clinical device utilized in this study provides sub-micron resolution images, enabling the detection of cellular metabolic changes via the TPEF signal emitted from cellular NADH. We performed a quantitative assessment using mitochondrial clustering analysis, based on the detection of the NADH TPEF signal (see *Methods*), on the MPM images obtained from the same graft throughout the treatment duration for each patient.

Before initiating treatment, we captured baseline data by imaging both the vitiligo area and the adjacent normal skin. The imaging biomarker represented by mitochondrial clustering (β), indicated an altered metabolic state of keratinocytes across the epidermis of vitiligo lesions in comparison to the normal metabolic state of keratinocytes in perilesional skin for both groups of patients (responders and non-responders to treatment) (Figure 3A).

**Figure 3.**
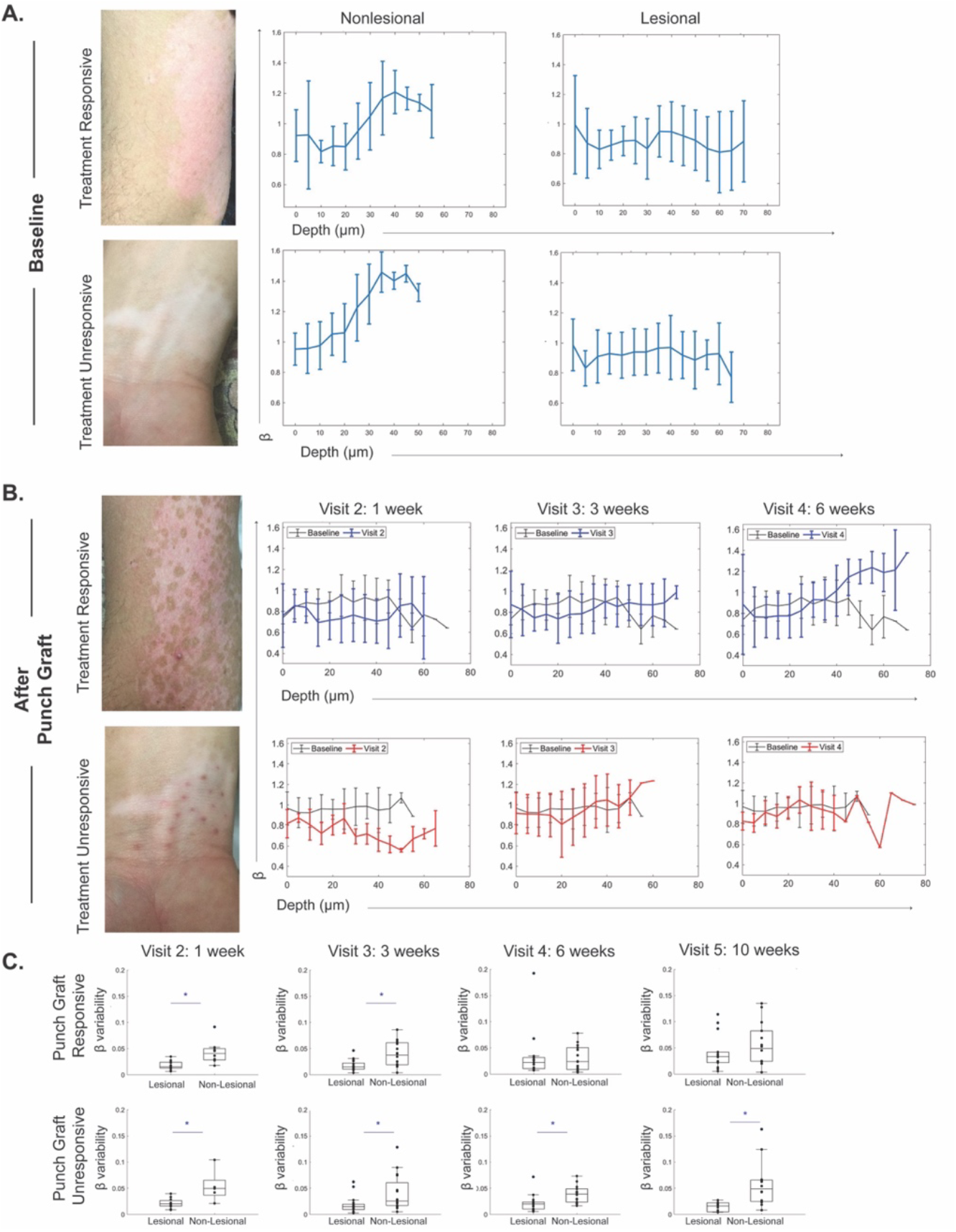
*In vivo* MPM imaging of vitiligo skin before and after punch grafting treatment show energy utilization shifts in treatment responders. **(A)** Representative baseline clinical images from vitiligo patients undergoing punch grafting in treatment responders and nonresponders. Average mitochondrial clustering (β) values based on z-stacks from 12 vitiligo lesions as a function of depth at baseline are shown as spline fits. **(B)** Patients underwent punch grafting and were imaged through 5 visits. Clinical images and spline fits at different visits were compared to baseline in responders (top) and nonresponders (bottom). β values for all lesions are shown. **(C)** Distribution of β variability values in punch grafting responders and nonresponders (n=12); each value corresponds to a z-stack of images acquired. * = p-value < 0.05.

Upon comparing the same metric at different treatment visits following punch grafting and nbUVB treatment, we found that basal keratinocytes in vitiligo skin that responded to treatment reverted to a more glycolytic metabolic profile within 6 weeks. In contrast, the basal keratinocytes in nonresponsive lesions maintained a metabolic profile, differing from that of adjacent normal skin (Figure 3B).

We evaluated the variability of the mitochondrial clustering (β) across the epidermis of vitiligo skin in responsive and nonresponsive cases and found that the metrics became similar to nonlesional skin in treatment responders by 6 weeks and remained consistent after 10 weeks (Figure 3C). Essentially, the metabolic profile differences between keratinocytes in vitiligo lesional and nonlesional skin were no longer significant in treatment responders at 6 weeks, while these differences persisted in treatment non-responders beyond 10 weeks after punch grafting.

## Discussion

In this study, we have demonstrated the capabilities of RCM and MPM imaging technologies in non-invasively monitor the cellular treatment response in vitiligo patients. Traditionally, measuring treatment response in single vitiligo lesions has been challenging. To date, there is no standardized measurement for assessing repigmentation in vitiligo[38]. The widely used clinical scoring system, known as the vitiligo area scoring index (VASI), primarily focuses on the percentage of affected body surface area (BSA) and the degree of depigmentation [39, 40]. However, VASI is inadequate for accurately capturing lesional repigmentation as it lacks sensitivity to detect small changes [41]. Thus, VASI changes after surgical procedures are typically evaluated 6-12 months after the procedure when more noticeable global changes occur [42]. Through analysis of RCM and MPM images, we have identified optical biomarkers that are associated with morohological changes in melanocytes and metabolic changes in keratinocytes occuring during punch-grafting and nbUVB treatment in stable vitiligo patients. These biomarkers provide a way of measuring repigmentation in individual lesions, which is particularly valuable as treatment strategies continue to advance in vitiligo patients. Our studies indicate that noninvasive imaging can directly detect the cytologic changes that correlate with therapy response and provides a way to directly measure how a given therapy can achieve melanocyte reconstitution. This direct measurement is superior to current clinical methods, where the quantification of response can be confounded by other factors (sun exposure, background skin pigmentation level, etc).

Through monitoring optical biomarkers associated with morphological changes in melanocytes derived from the RCM images over time, we made several significant observations. Firstly, we detected early melanocyte activation as soon as 3 weeks after treatment initiation, before clinical evidence of repigmentation. This activation was identified based on the presence of stellate dendrites, a characteristic feature of active melanocytes [32]. Moreover, we found that vitiligo patients who did not respond to treatment exhibited no melanocyte activation. This observation supports the hypothesis that repigmentation failure may be attributed to the arrest of outward melanocyte migration. This finding is consistent with a previous study using sequential immunohistochemistry analysis of vitiligo punch graft treatment, which demonstrated the absence of melanocytes in nonresponder tissue [43]. While these findings are promising, it is important to acknowledge the limitations imposed by the small sample size, necessitating further validation in future studies.

Our analysis of RCM images at different time points revealed that in the majority of treatment-responsive patients, the distance of melanocytes from the graft increased over time. This observation suggests melanocyte migration, although the limited sample size prevented us from determining the statistical significance of this result. Additionally, it should be noted that the melanocytes captured at different distances from the graft are likely not the same cells at different time points. Dynamic imaging would have provided further clarity; however, it was not feasible within the scope of this study due to time constraints for clinical imaging.

We gained additional insights by monitoring the optical biomarker associated with metabolic changes in keratinocytes over time, as derived from the MPM images. Prior to treatment, keratinocytes in vitiligo lesions exhibited an altered metabolic state, compared to the normal metabolic state of keratinocytes in perilesional skin, observed in both treatment-responder and non-responder groups. Subsequent analysis of the same biomarker at different treatment visits following punch grafting and nbUVB treatment revealed that basal keratinocytes in vitiligo skin responding to treatment underwent a metabolic shift towards a more glycolytic profile within 6 weeks. In contrast, nonresponsive lesions presented a sustained metabolic profile distinct from adjacent normal skin. These findings are in line with other studies that suggest keratinocyte-derived signals are important for melanocyte migration and build upon recent work [35, 44-46]. The results provide support for the hypothesis that the metabolic state of keratinocytes plays a role in the repigmentation of vitiligo, and failure to restore normal metabolism may serve as an indication of treatment response failure. From a practical point of view, these results support the development of newer grafting methods that require complete ablation of vitiligo skin and reconstitution of the graft site with keratinocytes and melanocytes from normal skin. Recently, fibroblast subsets have also been implicated in vitiligo pathogenesis [46, 47] but fibroblasts were not assessed in this study due to depth limitations of our imaging modalities. Future studies will examine whether spatial heterogeneity in grafting response observed in these methods is a result of heterogeneity in the metabolism of fibroblasts in grafted areas after transplantation.

While both RCM and MPM are capable of capturing active melanocytes, RCM had a distinct advantage in this study by providing a larger imaging area, enabling accurate monitoring of melanocyte migration throughout the treatment period. However, recent advancements in MPM technology, such as the development of the fast, large area multiphoton exoscope (FLAME) by our group [48, 49], hold promise for future studies. With devices like FLAME, it would be possible to capture changes in keratinocyte and melanocyte within the same field of view. This would provide valuable insights into how the interactions between melanocytes and keratinocytes evolve during the repigmentation process in both treatment responders and non-responders following punch grafting.

In conclusion, this study demonstrates the potential of RCM and MPM imaging technologies to non-invasively monitor cellular treatment response in vitiligo patients. Optical biomarkers associated with melanocyte morphological changes and keratinocyte metabolic shifts provide valuable insights into vitiligo persistence and early treatment response detection. Notably, our findings revealed early melanocyte activation as an early indicator of treatment response, observed as early as 3 weeks after treatment initiation, before clinical evidence of repigmentation. Additionally, the absence of melanocyte activation in non-responsive patients supports the notion that treatment failure may be attributed to the arrest of melanocytes from outward migration rather than immune-mediated destruction, supported by the observation that we also did not observe increased inflammation in any of the treated patients. However, it is essential to note that the small sample size emphasizes the need for further validation in future studies, and it is impossible to exclude the possibility of a small population of T cells not observed via imaging is preventing the initial egress of melanocytes from grafted skin. While RCM has proven advantageous in capturing melanocyte migration, the advancements in MPM technology offer opportunities to capture changes in both keratinocytes and melanocytes within the same field of view. These optical biomarkers also offer a novel approach to measure lesional repigmentation by a mechanism that directly visualizes the cytologic response, which can be incorporated into phase II studies whose logical outcome would be to measure the effect of therapies on melanocyte migration. Further validation is necessary to establish the reliability of these measurements before they can be implemented into a clinical trial workflow.

## Materials and methods

### Study Design

This study was conducted in accordance with the approved protocol from the Institutional Review Board (IRB 2018-4362) for clinical research in human subjects at UC Irvine, with the informed consent of each patient. Our study focused on investigating the effects of punch grafting and nbUVB treatment on vitiligo lesions using RCM and MPM imaging technologies.

Punch grafting is a surgical procedure involving the transplantation of 1 mm pieces of normal epidermis (commonly from the thigh area) into vitiligo-affected skin to establish a reservoir of melanocytes for repigmentation [50]. The procedure was performed by a board-certified dermatologist (AKG) and nbUVB therapy was initiated one week later. The inclusion criteria for stable vitiligo lesions required stability for at least one year and the absence of koebnerization, confetti-like depigmentation or trichome [51]. Patients were unresponsive to past treatment attempts, and had no treatment in the three months before imaging for this study.

Non-lesional sites on the thigh, exhibiting normal-appearance and absence of depigmentation under Wood’s lamp examination, were selected as the donor site for the punch grafting procedure. To ensure consistency, the area of interest after punch grafting was photographed using an iPad with a dermatoscope attachment, with anatomical landmarks serving as reference points for consistent imaging location.

A total of 11 patients with stable vitiligo were enrolled in this study with 12 lesions imaged. The vitiligo lesions were distributed across various locations, including wrist (2), hand (2), leg (5), arm (1), back (1) and face (1) (Table 1). Imaging sessions took place at baseline (before treatment initiation) and at 3, 6, and 10 weeks following the punch grafting and nbUVB treatment.

### RCM Imaging

We employed the Vivascope 1500 (CaliberID, MA, USA) for *in vivo* RCM imaging of vitiligo patients. This imaging system utilized an 830nm diode laser, a 30x objective with a numerical aperture of 0.9. The RCM images were obtained as z-stacks of en-face images, capturing the stratum corneum to the superficial dermis with a 5μm step size. Tile mosaics of en-face images were also acquired, covering areas of up to 6 × 6 mm. Each individual en-face image had a field-of-view of 500 × 500 μm^2^.

Due to the limited availability of the RCM device and participant dropout, RCM Imaging was not performed on all 11 enrolled vitiligo patients at every time point. Thus, imaging sessions took place at the 3-week and 6-week time point for 8 out of 12 lesions and at the 10-week time point for 11 out of the 12 lesions. Throughout the study, we ensured consistency by imaging the same graft and its surrounding area during each visit.

### MPM imaging

We used the MPM-based clinical tomograph (MPTflex, JenLab, GmbH, Germany) for *in vivo* imaging of vitiligo patients. This system incorporates a femtosecond laser (Mai Tai Ti:Sapphire oscillator, sub-100 fs, 80 MHz, tunable 690–1020 nm; Spectra-Physics), an articulated arm with near-infrared optics, and a beam scanning module. Two photomultiplier tube detectors were used to simultaneously detect two-photon excited fluorescence (TPEF) and second harmonic generation (SHG) signals. The excitation wavelength used in this study was 760 nm, with TPEF and SHG signals detected in the 410 to 650 nm and in the 385 to 405 nm range, respectively. We used a Zeiss objective (40x, 1.3 numerical aperture, oil immersion) for focusing the laser light into the tissue. The laser power ranged from 5 mW at the surface to 30 mW in the superficial dermis.

To acquire MPM data, we captured z-stacks of en-face images spanning from the stratum corneum to the superficial dermis. Each optical section had a field of view of 100 × 100 μm^2^ with a 5 μm step between sections. All 12 enrolled vitiligo lesions were imaged using MPM. During the initial visit, we imaged the vitiligo lesional area as well as a normally pigmented area on the upper thigh as control. The 512 × 512 pixel images were acquired at an approximate rate of 6 seconds per frame. MPM imaging of skin relies on the TPEF signal from NADH, FAD, keratin, melanin, and elastin fibers as well as the SHG signal from collagen [52, 53]. These images served as the basis for mitochondrial clustering analysis of keratinocytes.

### Mitochondrial Clustering Analysis

Mitochondrial clustering was quantified using a previously described Fourier-based approach in which the intensity variations of a TPEF image are attributed to variations in mitochondrial NADH binding[54]. This stems from the fact that the NADH fluorescence efficiency is enhanced by 2 to 10-fold when it is bound to different co-enzymes in the mitochondria[55, 56]. Previous work has demonstrated that mitochondrial clustering analysis is sensitive to dynamic changes in mitochondrial fusion and fission, which is dictated by the metabolic demands of a cell[57]. Specifically, this approach was validated by demonstrating that dynamic changes in mitochondrial clustering of human skin epithelia confined to the basal layer in response to hypoxia were consistent with an expected enhancement in the relative levels of glycolysis [33]. In this work, mitochondrial clustering, or β, was extracted for each image in the same manner from analysis of the intensity variations isolated from the cellular cytoplasm. Cellular cytoplasm was segmented in a manner consistent with methods described in [35]. In brief, autofluorescence contributions from keratin, melanin, nuclei, interstitial space, and dermis (as identified via collagen second harmonic generation) were masked[35]. A SHG mask was primarily created to remove contributions from collagen and stromal autofluorescence at the interface of the epidermis and dermis. Contrast-limited adaptive histogram equalization (CLAHE) was applied to SHG images and features were subsequently segmented using Otsu’s global thresholding. The SHG mask was finalized by applying a median filter to remove noise and taking the complement of the image to mask features corresponding to the segmented signal. Features corresponding to highly autofluorescent biomolecules such as keratin and melanin were masked using similar methods. CLAHE was applied to TPEF images and an Otsu’s global threshold was calculated. Pixels with intensity values 1.5X greater than the Otsu’s global threshold were segmented and masked. This empirically determined threshold was applied to all optical sections and was determined based on the propensity to remove highly autofluorescent signatures without masking pixels from intermediate cell layers which would not contain fluorophores such as keratin and melanin. The removal of nuclear and interstitial regions was achieved by applying 3 serial bandpass filters to contrast-limited adaptive histogram equalized TPEF images. Remaining features were segmented using Otsu’s global thresholding. A circular mask with a 500-pixel diameter was created to remove dim image corner artifacts. Masks were finalized with the removal of objects less than 8-pixels in size. The final mask was applied to raw TPEF images. Remaining autofluorescent features were randomly cloned 5 times to fill a full field of view. The average power spectral density (PSD) of the 5 cloned images was fit with an equation of form R(k) = Ak^-β^ for spatial frequencies (k) corresponding to spatial scales less than 8.5-μm. The absolute value of the fitted exponent, β, represents the degree of mitochondrial clustering within the cytoplasm. The β variance (variability) across the depth of the epithelium based on analysis of the corresponding stack of images was reported. All steps were completed in MATLAB.

### Statistical Analysis

For RCM melanocyte number visualized per unit area quantification, a two-tailed student’s t-test was used to determine significance between clinical responders (mean = 4.8, standard deviation = 2.58) and clinical nonresponders (mean = 0, standard deviation = 0).

All statistical comparisons of β variability were made within the framework of a linear mixed effects model generated in SAS JMP Pro 14. Variables such as a patient’s responsiveness to treatment, imaging location (lesional or non-lesional), and visit number were modeled as fixed effects. Variables such as patient number and the z-stack number were modeled as random effects. Pairwise comparisons of lesional and non-lesional skin for responsive and unresponsive patients at particular visits were made using post hoc Tukey honest significant difference tests with the significance level set to α = 0.05.

### Conflicts of Interest

MB is coauthor of a patent owned by the University of California, Irvine (UCI), which is related to multiphoton microscopy (MPM) imaging technology. MB is cofounder of Infraderm, LLC, a startup spin off from the UCI, which develops MPM-based clinical imaging platforms for commercialization purposes. The Institutional Review Board and Conflict of Interest Office of the UCI, have reviewed patent disclosures and did not find any concerns. The rest of the authors state no conflict of interest.

## Data Availability Statement

No datasets were generated during this study.

## Acknowledgements

JS acknowledges the grant support from the NIH (5KL2TR1416-6). MB, AKG and IG acknowledge the grant support from NIBIB (R01EB026705). MB and AKG also acknowledge the grant support from NIAMS (R21AR073408) and from the Skin Biology Resource-Based Center, University of California, Irvine (P3011935106).

## Author Contributions Statement

Conceptualization: MB, AKG, IG

Data Curation: JS, GL, CP, PM

Formal Analysis: JS, GL, CP, PM, ME

Funding Acquisition: JS, MB, IG, AKG

Investigation: GL,CP

Methodology: MB, IG, AKG

Project administration: JS, GL, PM, AKG, IG, MB

Resources: JS, MB, AKG

Software: GL, CP

Supervision: IG, MB, AKG

Validation: JS, GL, CP, IG, AKG, MB

Visualiation: JS, GL, CP

Writing – original draft preparation: JS, GL, MB

Writing – review & editing: JS, GL, CP, IG, MB, AKG

